# LazypipeX: Customizable Virome Analysis Pipeline Enabling Fast and Sensitive Virus Discovery from NGS data

**DOI:** 10.1101/2025.04.29.651217

**Authors:** Ilya Weinstein, Olli Vapalahti, Ravi Kant, Teemu Smura

**Affiliations:** Department of Veterinary Biosciences, University of Helsinki, 00014 Helsinki, Finland; Department of Virology, University of Helsinki, 00014 Helsinki, Finland; Department of Virology and Immunology, University of Helsinki and Helsinki University Hospital, 00014 Helsinki, Finland; Department of Tropical Parasitology, Institute of Maritime and Tropical Medicine, Medical University of Gdansk, 81-519 Gdynia, Poland

**Keywords:** bioinformatics pipeline, metagenomics, mNGS, virus discovery, virome

## Abstract

Metagenomic next-generation sequencing (mNGS) is pivotal for detecting known and novel viruses in diverse samples; however, its efficacy relies on robust bioinformatics pipelines. We present LazypipeX with multiple advanced features, including flexible annotation strategies that can be adjusted to match different datasets and applications. Using a synthetic benchmark, we show that a simple two-round annotation strategy designed for speed and low false positive rate required in virus diagnostics can reduce execution time to a fraction of the BLASTN search without any loss in accuracy. Additionally, using real data, we show that annotation strategies based on combinations of fast-to-accurate homology searches reduce execution time from 5- to 20-fold compared to the BLASTN/BLASTP baseline. Using one of these “annotation chains”, we characterized multiple novel complete viral genomes that were missed by the Lazypipe v1/v2 and CZ ID analyses. LazypipeX is a highly efficient and versatile tool for virome analysis, offering customizable and transparent workflows that can facilitate virus discovery and identification in diverse mNGS applications.

## 1 INTRODUCTION

Rapid developments in mNGS have enabled virologists to assess the true diversity of viruses in various samples (1). This is particularly true for viruses that do not propagate in cell culture and for novel or emerging viruses that fail to be detected using conventional PCR- or antigen-based methods. mNGS is a high-throughput, unbiased technology that can identify both known and novel pathogens and has great potential for viral discovery, surveillance, and broad-spectrum clinical diagnostics (1–3).

Virome identification with mNGS is heavily dependent on bioinformatic analysis, and it is increasingly recognized that applied bioinformatic methods will significantly influence the results (3, 4). Bioinformatics analysis of mNGS data is an actively evolving field, with a wide range of tools available for individual steps and pipelines that support end-to-end workflows (1, 5). These pipelines can be categorized into three primary functions: characterizing viral diversity and abundance, virus identification or diagnostics, and facilitating novel virus discovery. In this study, we focused on pipelines for virus diagnostics and discovery. Open source pipelines that have been applied in virus diagnostics include metaMix (6), RIEMS (7), Kaiju (8), VIP (9), DAMIAN (2), Jovian (10), and CZ ID (3), with an excellent comparative evaluation presented by De Vries et al. (4). Pipelines for virus discovery focus on sensitivity and, as suggested by several authors, benefit from integrating protein alignments and/or protein domain annotation to identify highly divergent emerging and novel viruses, although at the cost of heavier computation (2, 3, 11, 12). Based on this notion, most pipelines that include protein-level alignments or protein domain annotations are applicable to virus discovery. There are also a number of open-source pipelines that were specifically designed to support the discovery of novel pathogens from mNGS data, such as VirusSeeker (11), DAMIAN (2), CZ ID (3) and Microseek (12). We have previously published two mNGS pipelines, Lazypipe1 and Lazypipe2, primarily designed for virus discovery (13, 14), which have been successfully applied to characterize viromes and identify novel viruses in clinical samples (15), ticks (16–18), mosquitoes (19, 20), snakes (18), farm animals (21, 22), and wildlife (23–27).

Here, we describe LazypipeX with multiple new features, including the novel concept of annotation strategies that can incorporate multiple nucleotide-, protein- and hidden Markov model (HMM)-based searches into contig annotations. Using a synthetic benchmark, we showed that a conceptually simple two-round strategy, where the first round uses a fast Minimap2 search (28) against a virus-only database, followed by a second round using BLASTN (29) against larger databases to filter false positives, can achieve a 4- to 8-fold speedup compared to a one-round BLASTN search, without any loss in accuracy. When combined with background filtering, the speedup achieved was 30-fold, with minimal loss in precision. Using real data, we showed that combining varying fast-to-accurate homology searches reduced the execution time by 5- to 20-times compared to the BLASTN/P baseline. Usability for virus discovery was illustrated by using LazypipeX to characterize multiple divergent novel viral genomes from arthropod data.

## 2 MATERIAL AND METHODS

### 2.1 LazypipeX outline

LazypipeX is a highly customizable UNIX-based pipeline for automated assembly and taxonomic profiling of metagenomic libraries, building upon the original Perl, C++, and R architectures to enable high-throughput virus identification and discovery. The workflow begins with quality preprocessing using fastp (30), followed by nontarget background read filtering using BWA-MEM (31), Sambamba (32), and SeqKit (33). Post-assembly via MEGAHIT (34) or SPADES (35), the pipeline introduces a modular “annotation strategy” concept, allowing users to arrange in chains (and rounds) multiple fast-to-accurate homology searches — such as Minimap2, BLASTN/P/X, DIAMOND, and HMMscan — against precompiled, regularly updated NCBI NT, UniRef100, and viral HMM databases (e.g., Pfam and NeoRdRp). Taxonomic binning is performed using a bitscore-weighted model to generate detailed abundance tables and interactive Krona graphs (36).

### 2.2 Improved background filtering

LazypipeX supports multiple background filters to remove host or contaminant eukaryotic reads and contigs. This pre-assembly and post-assembly filtering accelerates annotation and allows for a smaller reference database. Multiple filters can be applied in tandem for pooled samples (e.g. arthropods) or samples contaminated by multiple sources. Because human nucleic acids often contaminate libraries, filtering *Homo sapiens* along with other backgrounds is recommended. The pipeline includes precompiled background filters for ∼20 species, including human, mosquitoes, ticks, bats, and various mammals. Users can also add custom filters by providing a FASTA file path in the configuration file.

### 2.3 Flexible and transparent annotation strategies

LazypipeX introduces the concept of an “annotation strategy”, which can be tailored for different use cases. Users can select strategies that chain homology searches to improve sensitivity or strategies that arrange them in rounds to boost precision. We benchmarked several predefined one-round, two-round, and chained strategies for virus identification and discovery, all accessible via the pipeline command line.

#### 2.3.1 Fast one-round strategies

*Minimap.NT_Vi_ and Minimap.RefSeq_Vi_:* Minimap2 all contigs against NCBI NT or RefSeq viruses. These provide efficient preliminary annotations at the cost of higher false-positive rates.

#### 2.3.2 Two-round strategies for virus identification

Two-round strategies combine a fast initial search against a virus-only database (e.g., Minimap2) with a sensitive secondary search (BLASTN) against a broader database to prune false positives. This reduces the search space for the slower algorithm. For example, if the execution time scales linearly with the size of the query and reference data, the execution time for the annotation of all contigs with a slower search against the complete database can be expressed as:

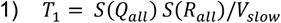

And the execution time for annotating only virus contigs in two-rounds can be expressed as:

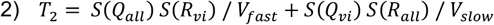

where *S(Q_all_)* is the size of the assembly, S(*Q*_*vi*_) is the size of the virus part of the assembly, *S(R_all_)* is the size of the reference database, *S(R*_*vi*_) is the size of the virus part of the reference, *V*_*slow*_ and *V*_*fast*_ are the speeds of the slower and faster search, respectively.

The ratio of execution times can be expressed as:

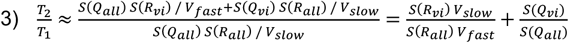

For example, if V_slow_/V_fast_ ≈ 10%, and if viruses make up ∼10% of the query data and 15% of the reference data, then T2/T1 ≈ 0.115, leading to a ∼9-fold speedup. We argue that these assumptions are realistic. For example, in the NT release from 2024/01/01, virus sequences constituted 15.3% of the database, and the proportion of assembled viral data in most metagenomic datasets is below 10% (e.g. in (37) library from rRNA depleted tissue RNA this was 1.68%). Furthermore, Minimap2, although not specifically evaluated against BLASTN, has been reported to be tens of times faster than long-read genomic mappers.

We tested three strategies using this logic:

- *Minimap.NT_Vi_ -Blastn.NT*_ABV_: In the first round, all contigs are mapped with Minimap2 against NT viruses (NT_Vi_; database descriptions are provided in Table 1). In the second round, pre-labelled viral contigs are annotated with BLASTN against NT archaea, bacteria, and viruses (NT_ABV_). This strategy includes background filtering and assumes that most eukaryotic sequences are removed before the annotation step. This allows to reduce the database for the second round to NT_ABV_.
- *Minimap.RefSeq_Vi_-Blastn.RefSeq_ABV_:* In the first round, all contigs are mapped with Minimap2 against RefSeq virus (RefSeq_Vi_). In the second round, pre-labelled viral contigs are annotated using BLASTN against RefSeq archaea, bacteria, and viruses (RefSeq_ABV_). This strategy also included background filtering.
- *Minimap.NT_Vi_-Blastn.NT:* In the first round, all contigs are mapped using Minimap2 against *NT*_*Vi*_. In the second round, the pre-labelled viral contigs are annotated using BLAST against the complete NT database.

**Table 1.** List of reference databases supported by LazypipeX.

| Database | Description |
| --- | --- |
| Minimap.NT<sub>ABV</sub>, Minimap.NT<sub>Vi</sub> | Fasta and accession-to-taxid files for annotations with Minimap2. NCBI NT archaea-bacteria-viruses and NT viruses. |
| Blastn.NT<sub>ABV</sub> and Blastn.NT<sub>Vi</sub> | BLASTN index for NT archaea-bacteria-viruses and NT viruses. |
| Minimap.RefSeq<sub>ABV</sub> and Minimap.RefSeq<sub>Vi</sub> | Fasta and accession-to-taxid files for annotations with Minimap2. RefSeq archaea-bacteria-viruses and RefSeq viruses. |
| Blastn.RefSeq<sub>ABV</sub> and Blastn.RefSeq<sub>Vi</sub> | BLASTN index for RefSeq archaea-bacteria-viruses and RefSeq viruses |
| Blastp.UniRef100<sub>ABV</sub> and Blastp.UniRef100<sub>Vi</sub> | BLASTP index for UniRef100 archaea-bacteria-viruses and viruses |
| Diamond.UniRef100<sub>ABV</sub> and Diamond.UniRef100<sub>Vi</sub> | Diamond index for UniRef100 archaea-bacteria-viruses and viruses |
| HMMscan.PFAM<sub>Vi</sub> | Viral HMM profiles from PFAM database (Mistry et al., 2021) |
| HMMscan.RdRp | NeoRdRp2 database (Sakaguchi et al., 2024) |

The first two require background filtering and utilize archaea-bacteria-virus (ABV) segments of NT and RefSeq. The third strategy omits filtering and uses complete NT, making it suitable for samples with complex backgrounds (e.g., fecal or environmental samples).

#### 2.3.3 Chained strategies for virus discovery

Chained annotation strategies maximize sensitivity by passing contigs with no hits through progressively more sensitive (and often slower) search engines. Strategies *Vi.chain1–4* require eukaryotic background filtering and utilize archaea-bacteria-virus (ABV) segments of NT and UniRef100. Vi.chain3.env omits filtering and uses complete databases, making it suitable for samples with complex backgrounds (e.g., fecal or environmental samples).

The chains are structured as follows (for database abbreviations see Table 1):

- *Vi.chain0* (Baseline): BLASTN (NT_ABV_) → BLASTP (UniRef100_ABV_)
- *Vi.chain1*: Minimap2 (NT_ABV_) → BLASTN (NT_ABV_) → BLASTP (UniRef100_ABV_)
- Vi.chain2: Minimap2 (NT_ABV_) → BLASTP (UniRef100_ABV_)
- Vi.chain3: Minimap2 (NT_ABV_) → DIAMOND blastp (UniRef100_ABV_)
- Vi.chain4: Minimap2 (NT_ABV_) → SANSparallel (UniProtKB)
- Vi.chain3.env: Minimap2 (NT) → DIAMOND blastp (UniRef100)

#### 2.3.4 Supported searches engines

These include Minimap2, BLASTN/P/X, DIAMOND blastp/x, SANSparallel, and HMMscan (28, 38–40). All searches utilized bitscore or e-value cut-offs defined in the main configuration file (default values given in Table 2). For protein-level searches (BLASTP, DIAMOND blastp, SANSparallel, HMMscan), ORFs ≥72 nt were extracted using MGA (default) or orfipy (41).

**Table 2.** LazypipeX default search engine parameters.

| Tool | Key Options | Filter/Selection |
| --- | --- | --- |
| Minimap2 | -x asm20 --secondary yes -cs -s 200 | peak_DP_alignment_score >= 200, top hits, no ties |
| BLASTN/P/X | -evalue 10 -max_target_seqs 5 | bitscore >= 200, top hits, no ties |
| DIAMOND blastp/x | --evalue 10 --max_target_seqs 5 --min-orf 72 --very-sensitive | bitscore >= 200, top hits, no ties |
| HMMscan | -E 0.01 | Top hits with ties |
| SANSParallel | -m SANStopHtaxid --SANS_H 5 -R | bitscore >= 200, top hits, no ties |

#### 2.3.5 Supported reference databases

These are listed in Table 1 and can be installed to local volumes using *install_db.pl* utility. The default nucleotide database is NCBI NT, which was also the most common reference database employed by eight out of 13 pipelines examined by de Vries et al. (4) and by CZ ID (3). For the default protein database, we chose UniRef100, similar to GenomeDetective, which uses UniRef90 (42). Other potential options include NCBI NR (e.g., DAMIAN (2) and CZ ID (3)) and RefSeq proteins (e.g., metaMix ((6)). However, we selected UniRef100 because it is highly representative, while remaining an order of magnitude smaller than the NR.

Supported databases were built from RefSeq (release 228), NT (version 2024/01/01), and UniRef100 (version 2024/11/27) by filtering archaeal, bacterial, and viral sequences using get_species_taxids.sh (43) and SeqKit (33), followed by default indexing calls. The Minimap2 database comprised FASTA files combined with accession-to-taxid TSV files. *HMMscan.PFAMVi* was compiled by downloading all viral HMM profiles from Pfam (44); we also used the original NeoRdRp 2.1 database (45).

### 2.4 Additional features

LazypipeX now integrates the ICTV VMR metadata (46), including *host.source* and *genome*.composition, into annotation tables. It also adds support for DIAMOND blastp/blastx (40) and HMMscan (47), with new ABV and Vi databases derived from UniRef100, Pfam (44) and NeoRdRp 2.1 (45).

LazypipeX can output IGV reports (48) for the identified viral contigs. The pipeline bins contigs by species, aligns them to the top reference hits via Minimap2, and produces interactive IGV visualizations of genome coverage and variants. These reports facilitate the identification of contig overlaps, sequence similarities, and gene coverage (e.g., polymerases) in fragmented assemblies. Example IGV reports for ten mosquito pools are available at https://doi.org/10.5281/zenodo.18348371.

### 2.5 Benchmarking

#### 2.5.1 Benchmarking background filtering

We evaluated LazypipeX background filtering on two datasets: a simulated human metagenome from MetaShot (49) and two *Rattus norvegicus* liver libraries (37). For the MetaShot benchmark, reads were filtered using a human reference genome (acc. GCF_000001405.40) and validated against the sequence labels included in the benchmark. For the rat samples, reads were filtered against the *R. norvegicus* reference (GCF_036323735.1) and validated by comparing results to LazypipeX BLASTN-based binning. Performance was assessed by iterating over the BWA MEM (31) alignment score (AS) and mapping quality (mapq) thresholds.

#### 2.5.2 Benchmarking strategies for virus identification on simulated data

We evaluated LazypipeX with one- and two-round strategies using a MetaShot simulated human metagenome (20.5M PE 150bp reads, 80 viral and 70 bacterial pathogens) (49).

To assess time savings versus accuracy, we compared two-round strategies (*Minimap.NT_Vi_-Blastn.NT_ABV_* and *Minimap.NT_Vi_-Blastn.NT*) against their one-round equivalents (*Blastn.NT_ABV_* and *Blastn.NT*). We also tested *Minimap.RefSeqVi-Blastn.RefSeq_ABV_*, standalone Minimap2 rounds (*Minimap.NT_Vi_, Minimap.RefSeq_Vi_*), and a common metagenomic approach, DIAMOND blastp against complete NR (*Diamondp.NR)*.

For performance baselines, we included Lazypipe v2.1 (Minimap against NT) and the CZ ID pipeline (Illumina v8.3, NCBI index dated 2024/02/06 with human host subtraction) (3). Most local runs utilized human background filtering (GCF_000001405.40). The databases used included RefSeq (rel. 228), NT (2024/01/01), UniRef100 (2024/11/27), and NR (2025/08/07). For all compared pipelines, reported taxids were linked to the NCBI taxonomy dated 2025/09/10 using a taxonkit (50). For CZ, ID read counts were selected from the NT count field when this was non-zero, and from the NR count field when the NT count was zero. For all pipelines, taxa with fewer than 10 reads and taxa labelled as bacteriophages were ignored. All local runs were performed on a Linux/Unix CPU supercluster with 30 cores, with each running at 2.1 GHz.

#### 2.5.3 Benchmarking virus discovery on real data

We benchmarked chained strategies (*Vi.chain0-4* and *Vi.chain3.env*) on Illumina libraries from 90 *Ochlerotatus* mosquito pools (acc. PRJNA852425) previously analyzed using Lazypipe1 and downstream BLASTX (19). Background was filtered using the human (GCF_000001405.40) and four mosquito reference genomes (GCF_002204515.2, GCF_035046485.1, GCF_016801865.2, and CA_964187845.1). The results were compared against *Vi.chain0* baseline and Lazypipe2.

A novel horsefly (*Tabanidae sp*.) library was used to test HMMscan integration. Filtering utilized a related *Chrysops caecutiens* reference (acc. GCA_963971475.1). The annotation chain followed *Vi.chain1* (Minimap2 → BLASTN → BLASTP), supplemented with HMMscan (Pfam → NeoRdRp v2.1).

Finally, LazypipeX was compared against CZ ID (v8.3) using the horsefly and two mosquito samples (SRR19859382–3). All local runs used default settings on a 30-core (2.1 GHz) Linux cluster.

## 3 RESULTS

### 3.1 Background filtering

The results of background filtering on the simulated data are presented in Table 3. Filtering by mapping quality (mapq) was less effective than filtering by alignment score (AS). At the default threshold (AS ≥ 100), 99.998% of human reads were removed, while all non-retroviral reads were retained. Notably, 96.375% of human endogenous retroviral (HERVs) reads were removed, likely due to the presence of these sequences in the reference genome. For accurate ERV binning, we recommend strategies that bypass background filtering (see Discussion). The results for the rat samples (Supp. Table 1) similarly favored AS filtering; the default threshold (AS ≥ 100) removed 63.1-70.2% of rodent reads while retaining 99.95-100.00% viral and ∼100.00% bacterial reads.

**Table 3.** Benchmarking background filtering on simulated data from (49). Selected filter highlighted. Filter, applied filter with AS representing alignment score and mapq mapping quality threshold, Total, total fraction of reads filtered, Homo sapiens, human reads filtered, Bacteria, bacterial reads filtered, Viruses, all viral reads filtered, HERV, reads from Human Endogeneous Retroviruses filtered, Viruses excl. Retroviridae, filtered viral reads excluding Retroviridae reads.

| Filter | Total | Homo sapiens | Bacteria | Viruses | HERV | Viruses excl. Retroviridae |
| --- | --- | --- | --- | --- | --- | --- |
| AS0 | 99.587 % | 100.000 % | 91.696 % | 97.529 % | 99.544 % | 96.804 % |
| AS30 | 94.687 % | 100.000 % | 0.021 % | 21.982 % | 96.453 % | 0.151 % |
| AS50 | 94.686 % | 99.999 % | 0.002 % | 21.913 % | 96.450 % | 0.046 % |
| AS75 | 94.685 % | 99.999 % | 0.001 % | 21.887 % | 96.444 % | 0.008 % |
| AS100 | 94.684 % | 99.998 % | 0.000 % | 21.861 % | 96.354 % | 0.000 % |
| AS125 | 94.681 % | 99.997 % | 0.000 % | 21.701 % | 95.649 % | 0.000 % |
| AS150 | 76.305 % | 80.631 % | 0.000 % | 11.793 % | 51.980 % | 0.000 % |
| mapq0 | 99.587 % | 100.000 % | 91.696 % | 97.529 % | 99.544 % | 96.804 % |
| mapq10 | 90.708 % | 94.938 % | 16.278 % | 26.474 % | 52.932 % | 20.282 % |
| mapq20 | 89.851 % | 94.855 % | 1.954 % | 12.879 % | 50.600 % | 2.122 % |
| mapq30 | 89.159 % | 94.219 % | 0.382 % | 10.595 % | 45.364 % | 0.459 % |
| mapq40 | 89.043 % | 94.117 % | 0.038 % | 10.147 % | 44.656 % | 0.022 % |
| mapq50 | 88.564 % | 93.623 % | 0.005 % | 8.680 % | 38.240 % | 0.005 % |
| mapq60 | 88.285 % | 93.331 % | 0.001 % | 8.266 % | 36.431 % | 0.000 % |

### 3.2 Virus identification from simulated data

We evaluated LazypipeX (several strategies), Lazypipe2, and CZ ID using a human-simulated metagenome (49). The species-level precision, recall, F1-score (harmonic mean of precision and recall), execution time, memory usage, and erroneous predictions are detailed in Table 4.

**Table 4.**
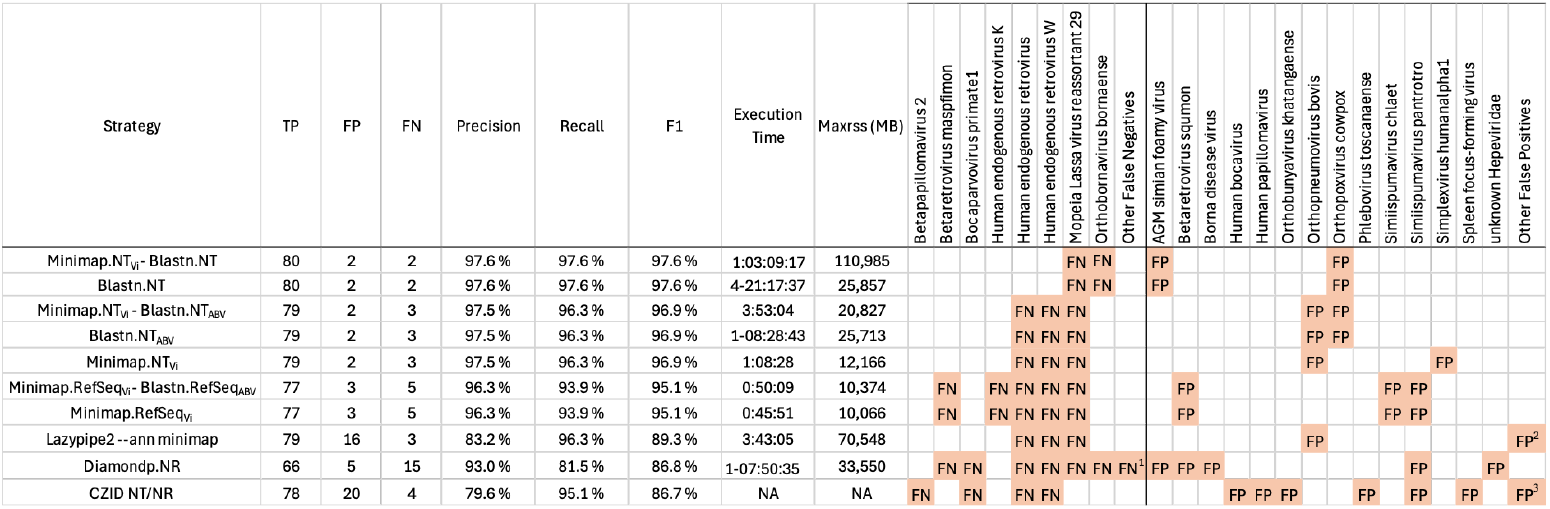
Accessing LazypipeX accuracy of virus taxon retrieval, execution time and memory consumption on simulated data from (Fosso et al., 2017). Compared pipelines are ordered by the descending F1-score. Column headers: TP, number of true positive species, FP, number of false positive species, FN, number of false negative species, F1, F1-score, Maxrss, maximum memory consumption. ^1^Other false negatives for Diamond.NR: Alphainfluenzavirus influenzae, Cytomegalovirus humanbeta5, Morbillivirus hominis, Orthoebolavirus zairense, Orthohepadnavirus hominoidei, Orthomarburgvirus marburgense, Phlebovirus napoliense, TTV-like mini virus and Uukuvirus uukuniemiense. ^2^Other false positives for Lazypipe2: Bacteriophage sp., Bifidobacterium phage Bbif-1, Caudoviricetes sp., Escherichia phage Lambda_ev058 and phi458, Herelleviridae sp., Inoviridae sp., Maltschvirus maltsch, Myoviridae sp. ctk251 and ctZhz2, RockefellervirusIPLA5, Siphoviridae sp. ctX581, Streptococcus phage IPP16, Streptococcus satellite phage Javan375 and uncultured human fecal virus. ^3^Other false positives for CZID: Biseptimavirus BU01, Caudoviricetes sp., Crassvirales sp., Myoviridae sp. ctk251/ctuJM17, Phietavirus B236, Podoviridae sp. ct7Ex2, Siphoviridae sp. ctCCX1/ctGz830/ctSqC25/ctX581/ctxvK3, uncultured human fecal virus and virus sp. ctJrn16.

The highest accuracy was achieved by LazypipeX annotations against the complete NT database, without host filtering. Both one-round (*Blastn.NT*) and two-round (*Minimap.NT_Vi_-Blastn.NT*) strategies yielded identical errors: two false negatives (FN) – *ML29* (misclassified as *Mammarenavirus lassaense*) and *Orthobornavirus bornaense* (contig unannotated), and two false positives (FP) – *Orthopoxvirus cowpox* (misclassified *Vaccinia virus*) and *ASM Simian foamy virus* (misclassified *Human spumaretrovirus*). The two-round version achieved a 4.3X speedup (∼27h vs. ∼117h).

LazypipeX annotations against *NT*_*Vi*_/*NT*_*ABV*_ (with background filtering) reported two FPs and three FNs – two filtered endogenous retroviruses and *ML29*. The execution time dropped from ∼32h (one-round) to ∼4h (two-round), an 8.4X speedup. LazypipeX annotations against the smaller RefSeq database (*Minimap.RefSeq_Vi_-Blastn.RefSeq_ABV_* and *Minima.RefSeq_Vi_*) had higher error rates, resulting in three FPs and six FNs. Overall, transitioning from one-round *BLASTN.NT* to two-round *NT*_*ABV*_ annotations (*Minimap.NT*_*Vi*_-*Blastn.NT*_*ABV*_) provided ∼30X speedup with identical recall and minimal precision loss.

Annotating with DIAMOND Blastp against NR (*Diamondp.NR*) showed low accuracy reporting five FPs and failed to detect 15 viral pathogens (Table 4). Lazypipe2 was hindered by low precision, primarily due to a high number of unlabeled bacteriophages. The CZ ID also reported multiple FPs, including four FP eukaryotic viruses and 14 unlabeled bacteriophages. The FNs for CZ ID included two retroviruses, likely filtered with the host genome, as well as *Betapapillomavirus 2* and *Bocaparvovirus primate1* (Table 4). The latter two were likely misclassified as *Human papillomavirus* and *Human bocavirus*, which may be attributed to discrepancies between the applied taxonomies.

### 3.3 Virus discovery from real data

#### 3.3.1 Benchmarking against BLAST baseline

We compared the identification of viral sequences from 90 mosquito samples (19) using five chained strategies (*Vi.chain1-4* and *Vi.chain3.env*) and Lazypipe2. As “true” viral sequences or taxa are unknown for real data, we used *Vi.chain0* ( BLASTN (NT_ABV_) → BLASTP (UniRef100_ABV_)) as the ground-truth baseline to measure the relative accuracy and time savings compared to BLAST-based workflows.

In this evaluation, we focused on viral sequences ≥ 500 bp, defining true positives (TP) as sequences identified by both the evaluated and baseline strategies, and false positives (FP) or negatives (FN) as sequences unique to the evaluated or baseline strategies, respectively. Precision, recall, and the F1-metric were calculated using the total base-pair counts for these categories (Fig. 1 and Supp. Table 2).

**Figure 1.**
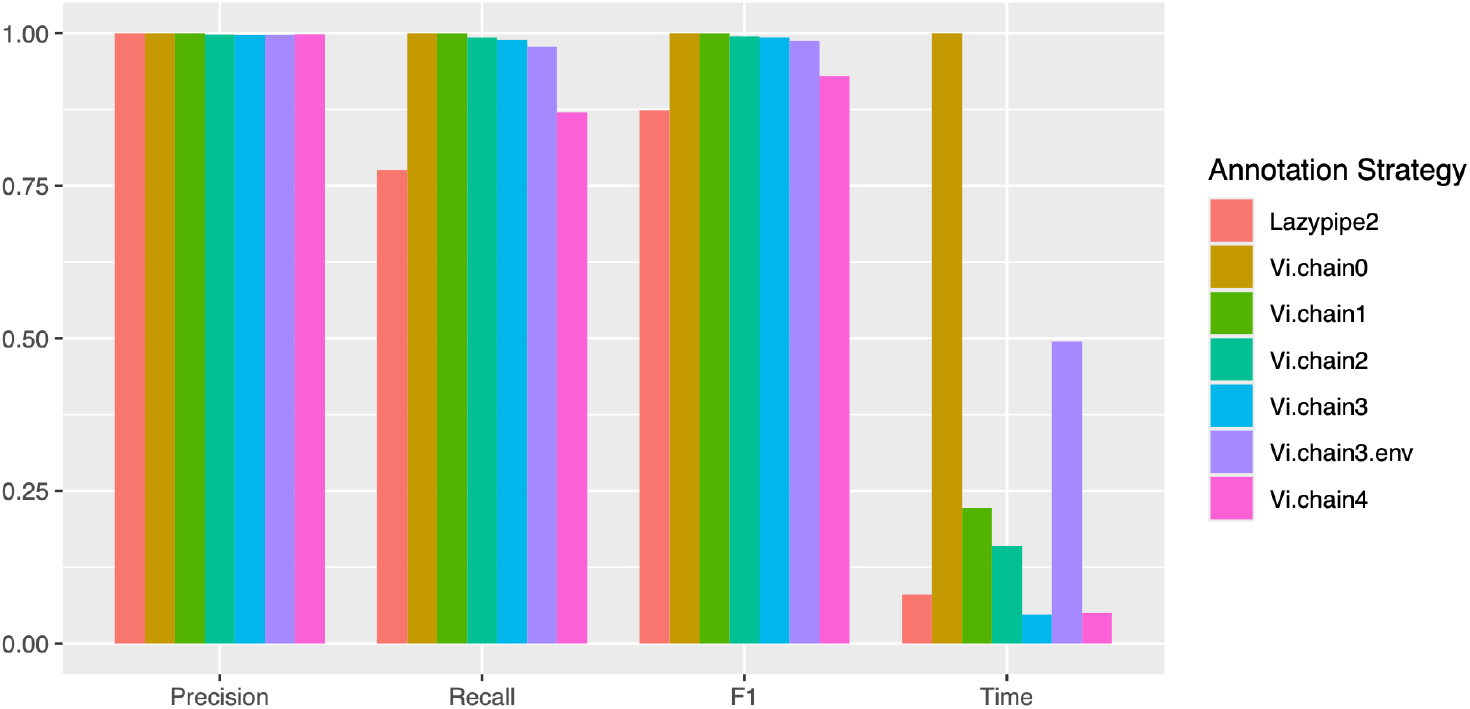
Overall performance of chained strategies (and Lazypipe2 –ann minimap) on the mosquito dataset. Precision, recall, F1-metric and Time were calculated relative to Vi.chain0 baseline as described in the text.

All LazypipeX strategies achieved ≥ 99.7% precision. *Vi.chain1* (introducing Minimap2 before BLASTN) provided a 5-fold speedup with negligible loss in recall (∼1 kbp of FNs). *Vi.chain2* and *Vi.chain3* (substituting BLASTN with Minimap2 and BLASTP with DIAMOND) yielded 6-fold and 20-fold speedups, respectively, with minimal loss in recall (∼31 kbp and ∼47 kbp of FNs, respectively). *Vi.chain3.env* (no background filtering) doubled the FNs to ∼91 kbp, likely due to contigs with close eukaryotic homologs excluded from the baseline. *Vi.chain4* (substituting BLASTP with SANSparallel) matched the 20-fold speedup of *Vi.chain3* but showed the lowest recall (87.0%). Lazypipe2 recall was significantly lower (77.6%) because it only permitted annotation against a single database. For this benchmark, the default NT database was used, which resulted in poor detection of distant homologs.

The average runtimes for mosquito samples (median library size 92.7 Mbp) were reduced from 20 hours to as little as 1 hour (*Vi.chain3/4*), illustrating time savings of up to 95% compared to the BLAST baseline.

#### 3.3.2 Identified viral sequences

Here, we characterized viral sequences identified using the LazypipeX *Vi.chain1* strategy (Minimap2 (NT_ABV_) → BLASTN (NT_ABV_) → BLASTP (UniRef100_ABV_)) from the mosquito dataset and compared these with previously published findings (19). We also examined a novel rhabdovirus discovered in a horsefly sample using *Vi.chain1* augmented with the HMMscan search. Viral discoveries were compared between *Vi.chain1* and CZ ID for the horsefly and two mosquito samples (SRR19859382-83). The detailed LazypipeX contig lists and CZ ID reports are provided in Supp. Table 3 and https://doi.org/10.5281/zenodo.18348611, respectively.

For the mosquito dataset, *Vi.chain1* identified 1,242 viral contigs (≥ 1 kbp; 3,559 kbp total). Of these, Minimap2 identified 78.1 %, BLASTN 1.6 % and BLASTP 20.3 %. To evaluate chained annotations for virus discovery, we focused on contigs identified via Minimap2 and BLASTP.

##### 3.3.2.1 Identified rhabdoviruses

Within *Vi.chain1*, Minimap2 identified 44 contigs (≥ 1 kbp; Supp. Table 3), covering all rhabdoviruses previously reported in this dataset (19). BLASTP identified two additional contigs representing complete rhabdovirus genomes, provisionally named *Kuusamo primrhavirus* and *Inari merhavirus*, whereas HMMscan detected a 12.7 kbp contig provisionally named *Lieksa stangrhavirus* (horsefly sample). Taxonomic classification and virus names were based on phylogenetic and ORF analyses (Supp. Text 1), incorporating the names of the Finnish municipalities where the samples originated.

##### 3.3.2.2 Identified negeviruses

Within *Vi.chain1*, Minimap2 identified 56 contigs (≥ 1 kbp; Supp. Table 3), covering all negeviruses previously reported for this dataset (19). BLASTP identified four additional complete negevirus-like sequences (11.1-11.4 kbp) and 21 partial sequences. Comparison of these BLASTP findings with previously published negeviruses (19) confirmed a very low nucleotide (nt) level homology (qcov ≤ 3%). The seven largest BLASTP findings were provisionally named as isolates of *Kuusamo* and *Hanko negev-like viruses* based on phylogenetic and ORF analyses (Supp. Text 1), and their municipalities of origin.

##### 3.3.2.3 Identified totiviruses and artiviruses

Within *Vi.chain1*, Minimap2 identified 315 contigs (≥ 1 kbp; Supp. Table 3), covering 49 of the 52 *Totiviridae* species previously reported for this dataset (19). This search failed to report *Hanko totivirus 10 (2.7 kbp), Ilomantsi totivirus 3* (1.2 kbp), and *Karstula totivirus (1.1 kbp)*, potentially due to high similarity to other totivirus-like sequences. BLASTP identified five additional complete totivirus-like whole genomes (7.5-7.7 kbp), which were provisionally named as isolates of *Joensuu artivirus* based on phylogenetic and ORF analyses (Supp. Text 1), and their municipalities of origin. Comparison with previously reported totivirus sequences (19) confirmed a very low nucleotide-level homology (qcov <14%) for all five contigs.

##### 3.3.2.4 Identified flaviviruses

Minimap2 identified 10 contigs (≥ 1 kbp; Supp. Table 3), covering all *Flaviviridae* species previously reported for this dataset (19). BLASTP identified six additional contigs that were mapped to *Flaviviridae*. These were provisionally named *Palkane flavivirus, Palkane-like flavivirus, Hameenlinna flavivirus*, and *Lestijarvi flavivirus* based on phylogenetic analysis (Supp. Text 1) and their municipalities of origin.

Comparison with previously reported *Flaviviridae* sequences (19) revealed 72.0% nucleotide identity between *Hameenlinna flavivirus* (acc. BK071037) and *Hanko virus* (acc. ON949929.1). All other sequences showed no significant homology at the nucleotide level.

#### 3.3.3 Comparison of virus discovery by CZ ID and LazypipeX

Comparing the CZ ID and LazypipeX findings for *Rhabdoviridae* from the horsefly and the two mosquito samples (acc. SRR19859382-83) revieled that, while CZ ID identified the complete *Inari merhavirus* genome, it only recovered the L-gene for *Lieksa stangrhavirus* and 138-606 bp fragments of *Kuusamo primrhavirus* (https://doi.org/10.5281/zenodo.18348611), all of which were captured as complete genomes using LazypipeX. Similarly, for negeviruses and arti- or totiviruses, CZ ID recovered fragmented sequences (0.1-1.1 kbp) and failed to report the full genomes of *Kuusamo negev-like virus* (isolate *FIN/PP-2018/83*) and *Joensuu artivirus* (isolate *FIN/PP-2018/83*) reported by LazypipeX. No *Flaviviridae* contigs were detected in either pipeline in these samples. Notably, CZ ID did not report any novel large (≥ 1 kbp) rhabdo, nege, arti, or totivirus sequences that were not captured by LazypipeX.

## 4. DISCUSSION

LazypipeX is a versatile and efficient pipeline for virus discovery and identification from metagenomic next-generation sequencing (mNGS) data, offering significant advancements in speed, accuracy, and flexibility.

The pipeline’s background filtering enables transparent removal of host and contaminant sequences by specifying database accessions and metadata in the configuration file. This approach helped achieve a marked speedup for a simulated benchmark with high host-sequence backgrounds (Table 4), driven by reduced data size and a smaller reference database for annotation. Benchmarking on real and synthetic datasets demonstrated high filtering accuracy (Table 3 and Supp. Table 1). The pipeline supports pre-built filters for common use cases, custom filter creation, and the simultaneous application of multiple filters. These capabilities make LazypipeX adaptable to diverse scenarios, such as clinical samples from human or animal patients, vector pools (e.g., mosquitoes and ticks), and wildlife specimens lacking species-level identification.

Like its predecessors, LazypipeX employs *de novo* assembly to generate contigs, enhancing the sensitivity and specificity of homology searches. This approach is shared by pipelines that classify assemblies instead of or in addition to classifying raw reads (2, 3, 6, 7, 10). LazypipeX integrates both nucleotide and protein alignments, a feature that is rare in mNGS pipelines. A recent comparison found that only two of the 13 pipelines used both alignment types (Alawi et al., 2019; Vilsker et al., 2019), and only DAMIAN incorporated Hidden Markov Models (de Vries et al., 2021). LazypipeX users can tailor the annotation procedure by chaining homology searches (e.g., Minimap2, BLASTN/P/X, DIAMOND, SANSparallel, and HMMscan) against different databases, optimizing the sensitivity or specificity depending on the dataset. This flexibility supports iterative adjustment of the pipeline to match diverse goals in viral metagenomics.

The two-round annotation strategy in LazypipeX significantly enhanced virus identification efficiency. An initial fast search against a virus-only database, followed by a second round using BLASTN against a larger database, achieved a four- to 8-fold speedup without compromising accuracy (Table 4). When combined with eukaryotic background filtering, this approach yielded a 30-fold speedup compared to single-round BLASTN against the complete NT database. Compared to Lazypipe2, DIAMOND blastp on NR, and CZ ID, LazypipeX’s two-round strategy (*Minimap.NT_Vi_-Blastn.NT* and *Minimap.NT_Vi_-Blastn.NT_ABV_*) demonstrated superior accuracy in identifying eukaryotic viral taxa (Table 4). For example, the DIAMOND blastp missed 15 viral pathogens, likely due to high protein-level similarity among related viruses. These results highlight the strength of LazypipeX’s nucleotide-based searches for the detection of known viruses.

LazypipeX two-round strategy is conceptually similar to other efforts to perform fast alignments prior to more accurate but slower refinement steps. The idea was inspired by the VirusSeeker pipeline, which starts with the fast identification of candidate viral sequences against virus-only nucleotide and protein databases, and then removes false positives with alignments against NCBI NT and NR (11). Similarly, in CZ ID, reads are first mapped with fast nucleotide and protein aligners prior to aligning contigs using BLAST against targeted databases (3). In GenomeDetective, viral reads are identified using a fast DIAMOND search on UniRef90 prior to assembling and running BLASTN/BLASTX on RefSeq (42). Lazypipe complements these approaches by offering the flexibility to choose different search engines and databases for both scanning and refinement stages.

LazypipeX’s chained annotation strategies proved to be highly effective for virus discovery. In a real dataset of 90 mosquito pooled samples, the *Vi.chain1* strategy (Minimap2/BLASTN/BLASTP) reduced execution time 5-fold compared to the BLASTN/P baseline, with precision and recall above 99.7% and 98.9%, respectively. Substituting BLASTP with faster tools, like DIAMOND or SANSparallel (*Vi.chain3, Vi.chain4*), achieved a 20-fold speedup, although *Vi.chain4* showed lower recall (87.0%). *Vi.chain1* retrieved nearly all previously reported viral sequences (19) via its initial Minimap2 search, while the subsequent BLASTP search identified novel divergent viruses, including multiple complete genomes. The older Lazypipe (v2), which is limited to one search at a time, had a much lower recall (78.6%). Incorporating HMMscan into *Vi.chain1* enabled the discovery of a novel rhabdovirus whole-genome sequence in a horsefly sample, which CZ ID only partially recovered (L gene only). Comparisons with CZ ID for rhabdo-, nege-, arti/toti-, and flaviviruses further demonstrated the superior recovery of novel genomes by lazypipeX. Findings by BLASTP in *Vi.chain1* were characterized using ORF, homology, and phylogenetic analyses, which confirmed that most of these were unrelated to viral sequences previously reported for this dataset, revealing complete genomes for at least six novel viruses (Supp. Text 1). These results underscore the utility of LazypipeX for detecting divergent viruses in complex mNGS data.

Based on our benchmarking, we proposed general guidelines for selecting LazypipeX strategies across various use cases (Table 5). For virus identification, we recommend two-round strategies against NT_ABV_ for tissue samples, while the complete NT database is preferable for samples potentially containing retroviral sequences or complex eukaryotic backgrounds (Table 5).

**Table 5.** Guidelines for selecting annotation strategies.

| Sequence Data | Analysis Goals | Recommended Annotation Strategy |
| --- | --- | --- |
| Illumina libraries from tissue samples and other samples with clearly defined eukaryotic background | Virus identification excluding retroviruses. BLASTN accuracy with up to 30X speedup compared to BLASTN on complete NT. | *background filter + Minimap.NT<sub>vi</sub> - Blastn.NT<sub>ABV</sub>* |
|  | Virus identification including retroviruses. BLASTN accuracy with up to 8X speedup compared to BLASTN. | *Minimap.NT<sub>vi</sub> - Blastn.NT* |
|  | Virus discovery excluding retroviruses. Precision/recall at ~100%/100% of combined BLASTN/BLASTP with up to 5X speedup. | *background filter + Vi.chain1* |
|  | Virus discovery excluding retroviruses. Precision/recall at ~100%/99% of combined BLASTN and BLASTP with up to 20X speedup. | *background filter + Vi.chain3* |
| Illumina libraries from environmental, fecal and other samples with complex eukaryotic backgrounds | Virus identification including retroviruses. BLASTN accuracy with up to 8X speedup compared to BLASTN. | *Minimap.NT<sub>vi</sub> - Blastn.NT* |
|  | Virus discovery including retroviruses. Precision/recall at ~100%/98% of combined BLASTN and BLASTP with up to 20X speedup. | *Vi.chain3.env* |

For virus discovery from tissue samples, we suggest using either *Vi.chain1* or *Vi.chain3. Vi.chain3* is particularly well suited for large datasets, as it achieved a 20-fold speedup relative to the BLASTN/P baseline with only a marginal reduction in accuracy (Supp. Table 2). Conversely, *Vi.chain1* offers a 5-fold speedup with virtually no loss in accuracy; this remains a viable option for analyzing smaller datasets or utilizing larger high-performance computing resources. For virus discovery from environmental, fecal, and other samples with complex eukaryotic backgrounds, we recommend using *Vi.chain3.env*, as this will also bin eukaryotic sequences.

## Future perspectives

We plan to extend the Lazypipe series to field-based applications, specifically targeting integration with the Oxford Nanopore MinION platform. Future iterations will be optimized for both offline and online processing modes, enabling rapid on-site analysis of mNGS data in remote or resource-limited settings.

## 4 Author Contributions

Conceptualization of the project: I.W., R.K., T.S., and O.V.; Conceptualization of the pipeline: I.W., R.K., and T.S.; Implementation, updates, and code distribution: I.W.; Reference Database Curation: I.W.; Benchmarking: I.W.; Testing and Reporting: I.W., R.K., and T.S.; Writing — Original Draft Preparation: I.W.; Writing — Review & Editing: O.V., R.K., and T.S.; Supervision of the study: R.K., T.S., and O.V.; Funding Acquisition: R.K. and O.V.

## 5 Acknowledgements

The authors wish to acknowledge the CSC IT Center for Science, Finland, for providing computational resources.

## 7 Data Availability Statement

The LazypipeX user manual and source code are available at https://bitbucket.org/plyusnin/lazypipe/ and https://www.helsinki.fi/en/projects/lazypipe. The virus sequences produced are publicly available in the NCBI GenBank database (accession codes provided in Supp. Table 4).

