## Supplementary material for "LazypipeX: Customizable Virome Analysis Pipeline Enabling Fast and Sensitive Virus Discovery from NGS data": Supp. Text 1

**Supplementary Text 1**

**Phylogenetic Analysis and Genetic Characterization for Selected Viral Findings**

### 1 Methods

#### 1.1 ORF analysis

ORF structure was visualized for selected viral findings using annotated array plots. Open reading frames (ORFs) were identified using orfipy (1) setting minimal ORF length to 300, excluding partial 5' ORFs, including partial 3' ORFs, and using the standard genetic code. Nested ORFs on either strand were excluded. Remaining ORFs were annotated with the top DIAMOND blastp (v2.1.10.164) hit (2) against the UniRef100 viral subset (database retrieved 27/11/2024). ORFs were grouped using reciprocal best BLASTP hits and manual curation, then labeled by the most frequent gene or protein name among annotations. Finally, plots were rendered using geneviewer (3).

#### 1.2 Phylogenetic analysis

Boostrapped maximum likelihood (ML) trees were constructed for selected viral findings and their references as follows. For rhabdoviruses (*Inari merhavirus*, *Kuusamo primrhavirus*, and *Lieksa stangrhavirus*), references included ICTV *Alpharhabdovirinae* (for *I. merhavirus*) or *Deltarhabdovirinae* (for the primrha- and stangrhaviruses), previously reported sequences for this dataset (4), and NCBI clustered NR homologs (BLASTX, bitscore ≥ 400). L-gene protein sequences ≥ 1,500 amino acids (AA) with confirmed BLASTP homology were retained.

For negevirus-like contigs, references were sourced from clustered NR homologs (BLASTX, bitscore ≥ 200, max 100 accessions) and *Negevirus* exemplar isolates as listed in (7). Protein sequences for the methyltransferase-helicase-RdRp region were extracted, excluding those < 1,000 AA or lacking BLASTP homology. Previously reported negeviruses (4) were omitted due to poor alignment with the new and reference sequences.

For *Artivirus*, references were sourced from clustered NR homologs (BLASTX, bitscore ≥ 200, max 100 accessions), all ICTV *Artitivirus* isolates, and previously published *Totiviridae* sequences (4). Protein sequences for both ORFs were extracted, excluding those with ORF1 < 1,000 AA or lacking BLASTP homology. Alignments for ORF1 and ORF2 were concatenated prior to tree construction.

For flavivirus contigs, references comprised clustered NR homologs (BLASTX, bitscore ≥ 200), ICTV Orthoflavivirus isolates, and previously reported Flaviviridae (4). Polyprotein sequences ≥ 1,000 AA with confirmed BLASTP homology were extracted for alignment.

All sequences were aligned with MAFFT v7.505 (5) (--retree 2 --maxiter 100) and trimmed using trimAl v1.2 (-automated1) (6). ML trees were inferred with IQ-TREE v2.1.3 (8) using 1,000 ultrafast bootstrap replicates and ModelFinder (9) to determine the best-fit substitution model. Trees were rooted at the midpoint, except for rhabdoviruses, which were rooted on *Infectious hematopoietic necrosis virus* (acc. L40883). Final trees were annotated in TreeViewer (10).

### 2 Results

#### 2.1 Novel rhabdoviruses

Bootstrapped ML trees were constructed from the L-gene for *Kuusamo primrhavirus*, *Lieksa stangrhavirus* and *Inari merhavirus.* In the phylogenetic tree (Fig 1), *Lieksa stangrhavirus* (acc. PV591081) formed a sister clade to a clade to five ICTV *Stangrhavirus* isolates, while *Kuusamo primrhavirus* (acc. BK070983) clustered within *Primrhavirus* genus, confirming the assignment of these sequences to the *Strangrhavirus* and *Primrhavirus* genera. *Inari merhavirus* (acc. BK070984) was positioned within the *Merhavirus* clade as an outgroup to the clade of Merida viruses (Fig 2), validating its assignment to the *Merhavirus* genus.

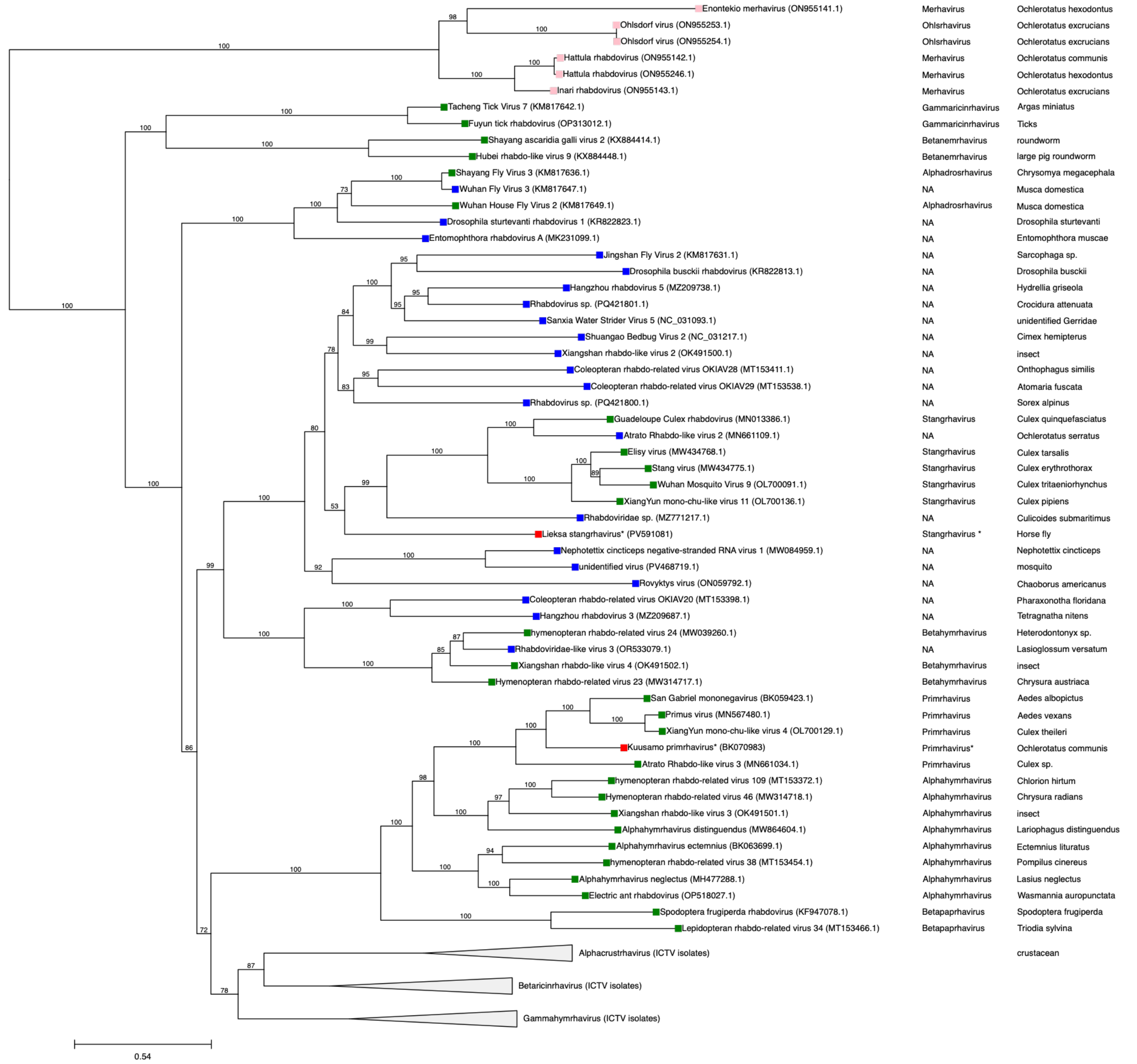

*Figure 1 Phylogeny of Lieksa stangrhavirus* and Kuusamo primrhavirus*. Tip colours: red, novel sequences, pink, Rhabdoviridae previously published for the mosquito dataset (4), green, ICTV Deltarhabdovirinae isolates, blue, homologs retrieved with BLASTX. Columns on the right denote virus genus and host species. Rooted on outgroup (infectious hematopoietic necrosis virus, acc. L40883, not shown), divergent clades collapsed for clarity. *Names assigned to new sequences are provisional.*

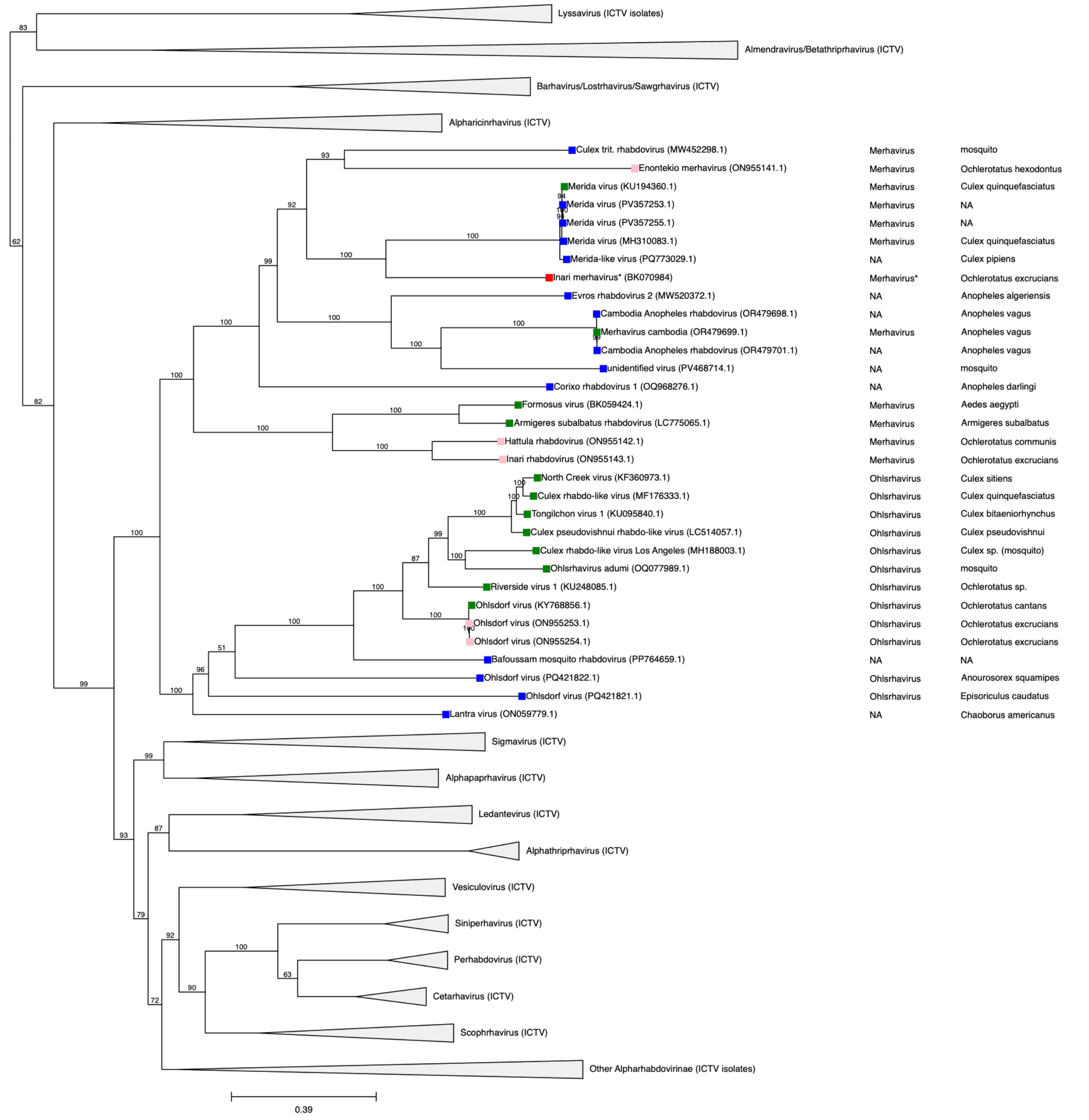


*Figure 2 Phylogeny of Inari merhavirus*. Tip colours: red, novel sequence, pink, rhabdoviruses previously identified from the mosquito dataset (4), green, ICTV Alpharhabdovirinae isolates, blue, homologs retrieved with BLASTX. Columns on the right denote virus genus and host species. Rooted on the outgroup (infectious hematopoietic necrosis virus, acc. L40883, not shown), divergent clades collapsed for clarity. *Names assigned to new sequences are provisional.*

Both *Kuusamo primrhavirus* and *Inari merhavirus* featured a five-ORF genome (N-P-M-G-L, Fig. 2) canonical for rhabdoviruses. In *Kuusamo primrhavirus*, ORFs 1, 2, 4 and 5 shared AA identities of 36.5%, 29.8%, 35.9%, and 52.3% with the nucleoprotein, phosphoprotein, glycoprotein, and RdRp of *Primus virus*, respectively. ORF3 lacked detectable homologs but is presumed to be the matrix (M) gene based on genomic position. In *Inari merhavirus*, ORFs 1, 3, 4 and 5 shared AA identities of

35.8%, 26.7%, 49.6%, and 52.1% to nucleoprotein, matrix protein, glycoprotein and RdRp in Merida-like viruses (accs. A0A894KPN5_9RHAB, A0A140DDD9_9RHAB, A0A5Q0TW57_9RHAB, A0A7L7QQ11_9RHAB), while ORF2 represents a putative phosphoprotein. Notably, the *Inari merhavirus* shared no significant nucleotide-level similarity with *Inari rhabdovirus* (acc. ON955143) previously reported for the same pooled sample.

*Lieksa stangrhavirus* exhibits a distinct architecture; ORFs 1, 4, 5, and 6 share homology with the nucleoprotein (24.7%), glycoprotein (24.4%), a secondary glycoprotein (21.9%), and the replicase (35.8%), respectively (accs. A0A0B5KRF8_9RHAB, A0A7D7F3C3_9RHAB, A0A8K1YQP7_9RHAB and A0A8K1YQP8_9RHAB). Although ORFs 2 and 3 lacked UniRef homologs, they likely represent the P and M genes.

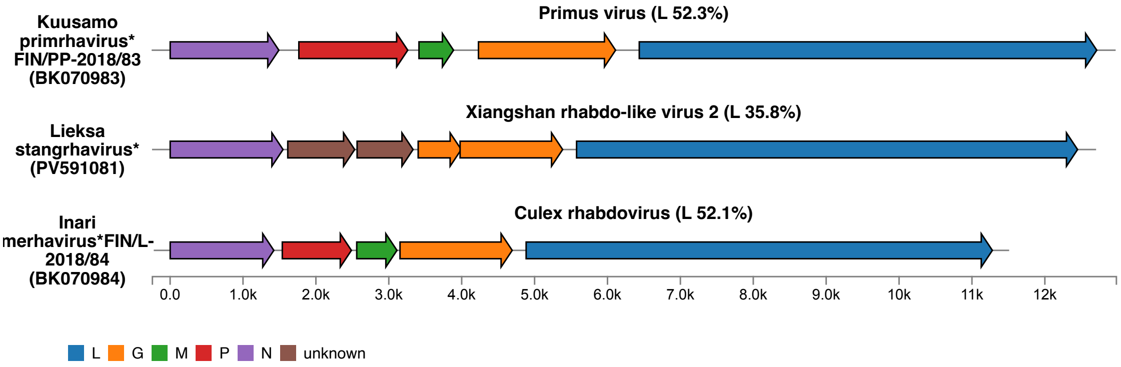


*Figure 2 ORF structure of novel Rhabdo-like sequences from mosquito samples and the horse-fly sample. Labels at the center represent closest homologs in UniRef with identity values in parenthesis. *Provisional virus names, isolates and accession numbers.*

Comparing *K. primhavirus* (acc. BK070983) protein sequences to ICTV *Primrhavirus* isolates (acc. MN661034, BK059423, MN567480, OL700129) showed above 66%, 46% and 63% amin acid (aa) divergence for the N, L, and G genes, respectively (Table 1), suggesting that this is a novel *Primrhavirus*. Unlike ICTV primrhaviruses, which were discovered in metagenomic studies of *Culex and Aedes* mosquitoes (11), *K. primrhavirus* was sequenced from *Ochlerotatus communis*, further supporting its demarcation as a separate species.

Comparing *L. stangrhavirus* (acc. PV591081) protein sequences to ICTV *Stangrhavirus* isolates (acc. MW434768, MN013386, MW434775, OL700091, OL700136) showed above 65%, 46% and 63% aa divergence for the N, L, and G genes, respectively (Table 1), suggesting that this is a novel *Stangrhavirus*. Additionally, the G gene in the new genome is split into two ORFs (Fig 2), and the host species is a horsefly, while ICTV stangrhaviruses are associated with mosquitoes.

Comparing *Inari merhavirus* (acc. BK070984) protein sequences to ICTV *Merhavirus* isolates (acc. OR479699, BK059424, ON955142, ON955143, KU194360, LC775065, AB604791) showed above 67%, 48% and 52% amino acid (aa) divergence for the N, L, and G genes (Table 1), suggesting that this is a novel *Merhavirus*. The aa divergence was similarly high (Table 1) when compared to *Merhavirus inari* (acc. ON955143), which was previously identified from the same sample. Thus, in the pooled sample, individual mosquitoes may have been infected or co-infected with two different merhaviruses.

*Table 1 Pairwise amino acid (AA) identity between the novel rhabdoviruses and ICTV isolates. Multiple sequence alignments were created with MAFFT for N, L and G-genes. P-value was then calculated in MEGA 12 discarding any pairwise gaps.*

| | | | AA Divergence* | | |
|---|---|---|---|---|---|
| Genus | Member Species | Accession | N gene | L gene | G gene |
| *Merhavirus* | ***Inari merhavirus*** | BK070984 | | | |
| | *M. cambodia* | OR479699 | 75.20 % | 57.10 % | 79.80 % |
| | *M. formosus* | BK059424 | 79.00 % | 60.90 % | 74.80 % |
| | *M. hattula* | ON955142 | 73.70 % | 61.10 % | 76.00 % |
| | *M. inari* | ON955143 | 74.80 % | 60.30 % | 76.00 % |
| | *M. merida* | KU194360 | 67.60 % | 48.20 % | 52.10 % |
| | *M. Subalbatus* | LC775065 | 75.80 % | 60.00 % | 75.40 % |
| | *M. tritaeniorhynchus* | AB604791 | 71.70 % | 55.80 % | 60.80 % |
| *Primrhavirus* | ***Kuusamo primrhavirus*** | BK070983 | | | |
| | *P. atrato* | MN661034 | 72.20 % | 51.40 % | 73.00 % |
| | *P. gabriel* | BK059423 | 66.00 % | 46.70 % | 63.10 % |
| | *P. primus* | MN567480 | 66.00 % | 47.60 % | 65.30 % |
| | *P. yunnan* | OL700129 | 65.80 % | 47.40 % | 65.60 % |
| *Stangrhavirus* | ***Lieksa stangrhavirus*** | PV591081 | | | |
| | *S. elisy* | MW434768 | 80.20 % | 67.60 % | 80.10 % |
| | *S. guadeloupe* | MN013386 | 79.10 % | 66.00 % | 75.70 % |
| | *S. stang* | MW434775 | 80.70 % | 66.70 % | 77.40 % |
| | *S. wuhan* | OL700091 | 81.80 % | 67.40 % | 74.20 % |
| | *S. yunnan* | OL700136 | 81.60 % | 68.00 % | 77.50 % |

#### 2.2 Novel Negev-like viruses

Bootstrapped ML tree was constructed using the RdRp gene for the seven largest negev-like virus sequences as described in the Methods (Fig 3). In the phylogenetic tree, the novel sequences were placed into two sister clades; isolates in the first clade were hosted by *Ochlerotatus caspius*, and in the second clade by *O. communis*. Both clades were distant from known members of the proposed genus *Negevirus* and clustered closest to negev-like viruses identified from *Bactrocera, Zeugodacus, Ceratitis*, and other *Diptera* (Fig 3). The ORF structure for these sequences is illustrated in Fig 4.

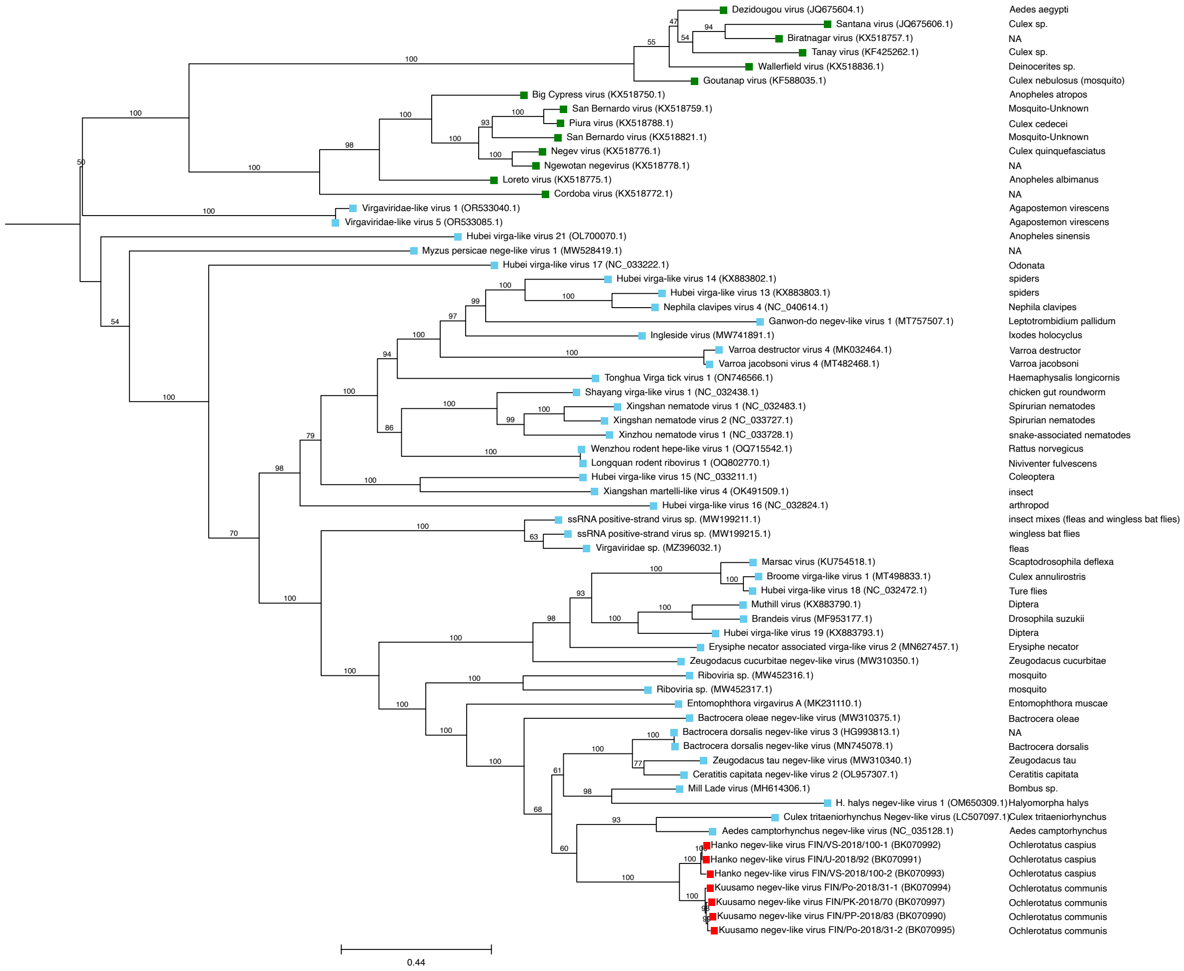


*Figure 3 Phylogeny for novel Negev-like virus sequences from mosquito samples. Tip colours: novel sequences in red, BLASTX homologs in blue, exemplar isolates of the proposed genus Negevirus (as listed in Nunes et al., 2017) in green. Host species in the right column. Scale bar is in Mya. Virus names assigned to new sequences are provisional.*

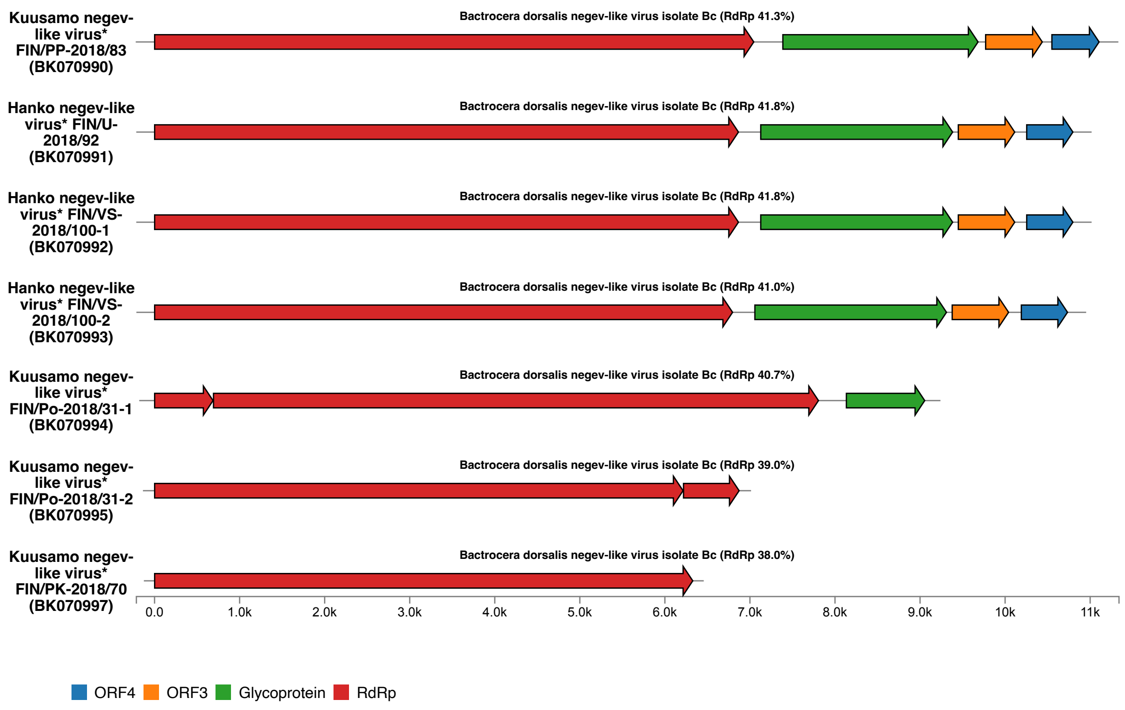


*Figure 4 ORF structure of novel Negev-like virus sequences from mosquito samples. Labels at the center represent closest homologs in UniRef with identity values in parenthesis. *Provisional virus names, isolates and accession numbers.*

Comparing the ORF1 protein sequence (concatenated methyltransferase, RNA-helicase, and RdRp) between the *Kuusamo negev-like virus* (acc. BK070997), *Hanko negev-like virus* (acc. BK070992) and exemplar isolates of the proposed genus *Negevirus* (as listed in (7)) showed <7% aa divergence in within-clade pairwise comparisons, >25% aa divergence in between-clade comparisons and >76% aa divergence to exemplar *Negevirus* isolates (Table 2). These divergence levels suggest that the seven isolates identified here likely represent two novel negevirus species hosted by *Ochlerotatus* mosquitoes.

*Table 2 P-value and amino acid (AA) identity within the two clades of novel Negev-like sequences and between clade representative isolates and exemplar species of proposed genus Negevirus. Multiple sequence alignments were created with MAFFT for concatenated protein sequences for Methyltransferase, RNA-helicase and RdRp. P-value was calculated in MEGA 12 discarding any pairwise gaps and converted to AA identity. Comparison was* *done between the representative variant (in bold) and all other sequences. Accessions for Negevirus exemplar species were collected from (Nunes et al., 2017). *Provisional virus names and isolates.*

| Group | Virus name | Accession | MTase-RNAhelicase-RdRp AA P-value | MTase-RNAhelicase-RdRp AA Identity |
|---|---|---|---|---|
| Novel Sequences | ***Kuusamo negev-like virus (FIN/PK-2018/70)**** | BK070997 | 0.000 | 100.00 % |
| | *Kuusamo negev-like virus (FIN/PP-2018/83)** | BK070990 | 0.066 | 93.37 % |
| | *Kuusamo negev-like virus (FIN/Po-2018/31-2)** | BK070995 | 0.065 | 93.53 % |
| | *Kuusamo negev-like virus (FIN/Po-2018/31-1)** | BK070994 | 0.014 | 98.62 % |
| | *Hanko negev-like virus FIN/VS-2018/100-1** | BK070992 | 0.282 | 71.79 % |
| | *Hanko negev-like virus FIN/VS-2018/100-2** | BK070993 | 0.290 | 71.04 % |
| | *Hanko negev-like virus FIN/U-2018/92** | BK070991 | 0.283 | 71.74 % |
| *Negevirus (Sandewavirus )* | *Dezidougou virus* | JQ675604 | 0.811 | 18.91 % |
| | *Santana virus* | JQ675606 | 0.815 | 18.53 % |
| | *Tanay virus* | KF425262 | 0.815 | 18.52 % |
| | *Goutanap virus* | KF588035 | 0.806 | 19.40 % |
| | *Biratnagar virus* | KX518757 | 0.809 | 19.07 % |
| | *Wallerfield virus* | KX518836 | 0.810 | 18.96 % |
| *Negevirus (Nelorpivirus)* | *Big Cypress virus* | KX518750 | 0.802 | 19.77 % |
| | *Brejeira virus* | KX518759 | 0.798 | 20.16 % |
| | *Corboda virus* | KX518772 | 0.810 | 19.03 % |
| | *Loreto virus* | KX518775 | 0.795 | 20.50 % |
| | *Negev virus* | KX518776 | 0.806 | 19.42 % |
| | *Ngewotan virus* | KX518778 | 0.801 | 19.91 % |
| | *Piura virus* | KX518788 | 0.803 | 19.71 % |
| | *San Bernardo virus* | KX518821 | 0.795 | 20.50 % |
| Novel Sequences | ***Hanko negev-like virus FIN/VS-2018/100-1**** | BK070992 | 0.000 | 100.00 % |
| | *Hanko negev-like virus FIN/VS-2018/100-2** | BK070993 | 0.058 | 94.20 % |
| | *Hanko negev-like virus FIN/U-2018/92** | BK070991 | 0.006 | 99.40 % |
| | *Kuusamo negev-like virus (FIN/PK-2018/70)** | BK070997 | 0.282 | 71.79 % |
| | *Kuusamo negev-like virus (FIN/PP-2018/83)** | BK070990 | 0.254 | 74.59 % |
| | *Kuusamo negev-like virus (FIN/Po-2018/31-2)** | BK070995 | 0.270 | 73.00 % |
| | *Kuusamo negev-like virus (FIN/Po-2018/31-1)** | BK070994 | 0.259 | 74.05 % |
| *Negevirus (Sandewavirus )* | *Dezidougou virus* | JQ675604 | 0.785 | 21.54 % |
| | *Santana virus* | JQ675606 | 0.786 | 21.41 % |
| | *Tanay virus* | KF425262 | 0.789 | 21.09 % |
| | *Goutanap virus* | KF588035 | 0.784 | 21.57 % |
| | *Biratnagar virus* | KX518757 | 0.789 | 21.08 % |
| | *Wallerfield virus* | KX518836 | 0.789 | 21.14 % |
| *Negevirus (Nelorpivirus)* | *Big Cypress virus* | KX518750 | 0.771 | 22.87 % |
| | *Brejeira virus* | KX518759 | 0.766 | 23.38 % |
| | *Corboda virus* | KX518772 | 0.779 | 22.09 % |
| | *Loreto virus* | KX518775 | 0.769 | 23.05 % |
| | *Negev virus* | KX518776 | 0.772 | 22.84 % |
| | *Ngewotan virus* | KX518778 | 0.768 | 23.21 % |
| | *Piura virus* | KX518788 | 0.765 | 23.52 % |
| | *San Bernardo virus* | KX518821 | 0.763 | 23.71 % |

#### 2.3 Novel Artivirus

BLASTP identified five complete totivirus-like whole genome contigs ranging from 7.5 to 7.7 kbp, with the ORF structure presented in Fig 4.

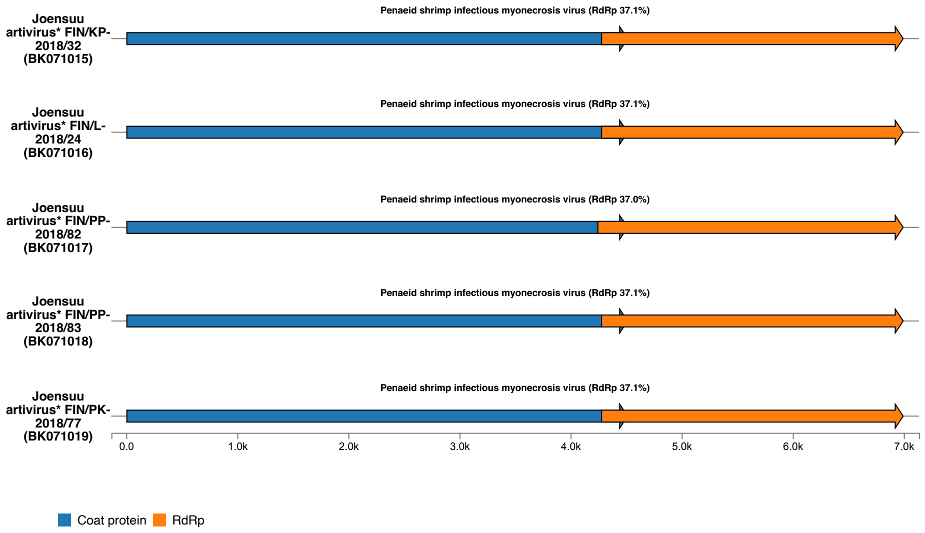


*Figure 4 ORF structure of novel Artivirus sequences from mosquito samples. Labels at the center represent closest homologs in UniRef with identity values in parenthesis. *Provisional virus names, isolates and accession numbers.*

Bootstrapped ML tree was constructed with concatenated ORF1 and ORF2 aa sequences. *Joensuu artivirus* formed a monophyletic sister clade to the unclassified *Grogonang virus 29* (OR270267) (Fig 5). The two closest classified viruses were *Artivirus sani* (acc. EU715328) and *Artivirus ni* (acc. GQ342961), confirming the assignment of the novel sequences to *Artivirus* (Fig 4).

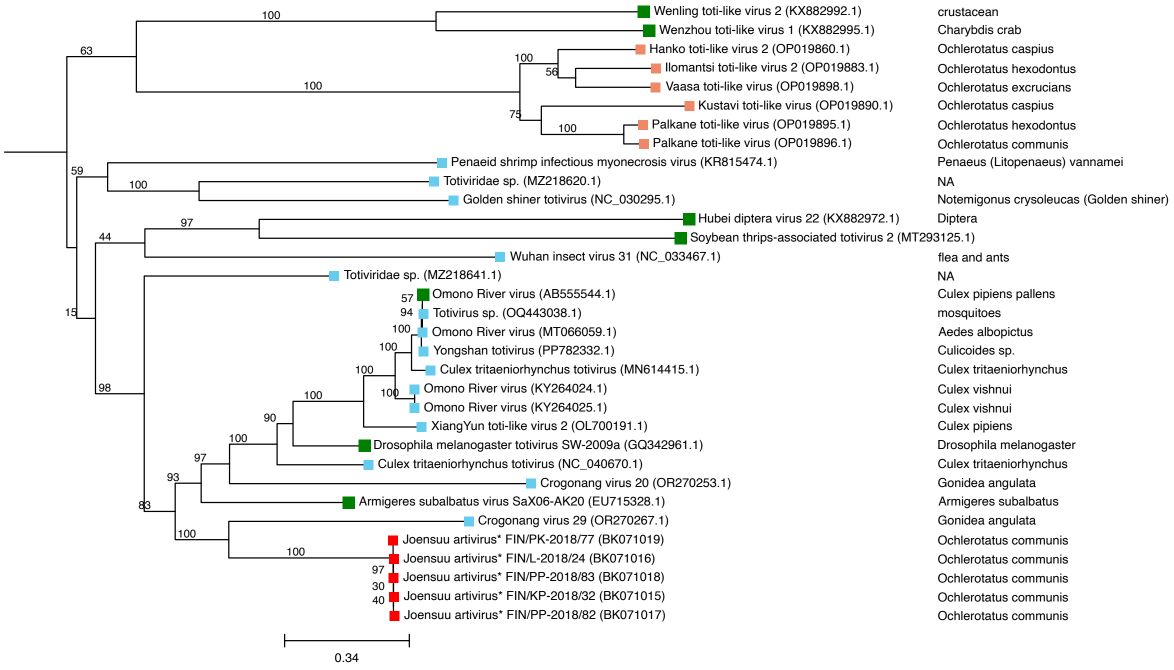


*Figure 5. Phylogeny of novel Artivirus sequences from mosquito samples. Tip colours: novel sequences in red, Totiviridae sequences previously published for this data in light red, BLASTX homologs in blue, ICTV Artivirus exemplars in green. Host species in denoted in the right column. Scale bar is in Mya. *Provisional virus names, isolates and accession numbers.*

*Joensuu artivirus FIN/PK-2018/77* (BK071019) was selected as the representative isolate, and its RdRp aa sequence was compared to other members of the clade and established member species of *Artivirus* (acc. KX882995, KX882989, AY570982, MT293125, GQ342961, KX882972, EU715328, AB555544, KX882992). The representative isolate shared <40% aa identity with established *Artivirus* species and >99% aa identity within other *Joensuu artivirus* sequences (Table 3), suggesting that these isolates represent a novel *Artivirus* species.

*Table 3 P-value and amino acid (AA) identity within the Joensuu artivirus clade and between the representative Joensuu artivirus isolate and ICTV Artivirus isolates. Multiple sequence alignments were created with MAFFT for RdRp. P-value was calculated in MEGA 12 discarding any pairwise gaps and converted to AA identity.*

| Group | Virus name | Accession | RdRp AA P-value | RdRp AA Identity |
|---|---|---|---|---|
| Novel Sequences | ***Joensuu artivirus (FIN/PK-2018/77)**** | BK071019 | | |
| | *Joensuu artivirus (FIN/L-2018/24)** | BK071016 | 0.00442 | 99.56 % |
| | *Joensuu artivirus (FIN/PP-2018/83)** | BK071018 | 0.00552 | 99.45 % |
| | *Joensuu artivirus (FIN/KP-2018/32)** | BK071015 | 0.00552 | 99.45 % |
| | *Joensuu artivirus (FIN/PP-2018/82)** | BK071017 | 0.00552 | 99.45 % |
| *ICTV Isolates* | *Artivirus go* | KX882995 | 0.66626 | 33.37 % |
| | *Artivirus hachi* | KX882989 | 0.66667 | 33.33 % |
| | *Artivirus ichi* | AY570982 | 0.6228 | 37.72 % |
| | *Artivirus kyu* | MT293125 | 0.68415 | 31.59 % |
| | *Artivirus ni* | GQ342961 | 0.60813 | 39.19 % |
| | *Artivirus roku* | KX882972 | 0.70138 | 29.86 % |
| | *Artivirus sani* | EU715328 | 0.61879 | 38.12 % |
| | *Artivirus shi* | AB555544 | 0.62663 | 37.34 % |
| | *Artivirus shichi* | KX882992 | 0.65934 | 34.07 % |

#### 2.4 Novel flaviviruses

BLASTP in *Vi.chain1* identified six contigs mapping to *Flaviviridae* with an ORF structure presented in Fig 5. All six contigs contained long ORFs, complete at the 5' end and partial at the 3' end, with flavivirus polyprotein homologs in UniRef (Fig 5).

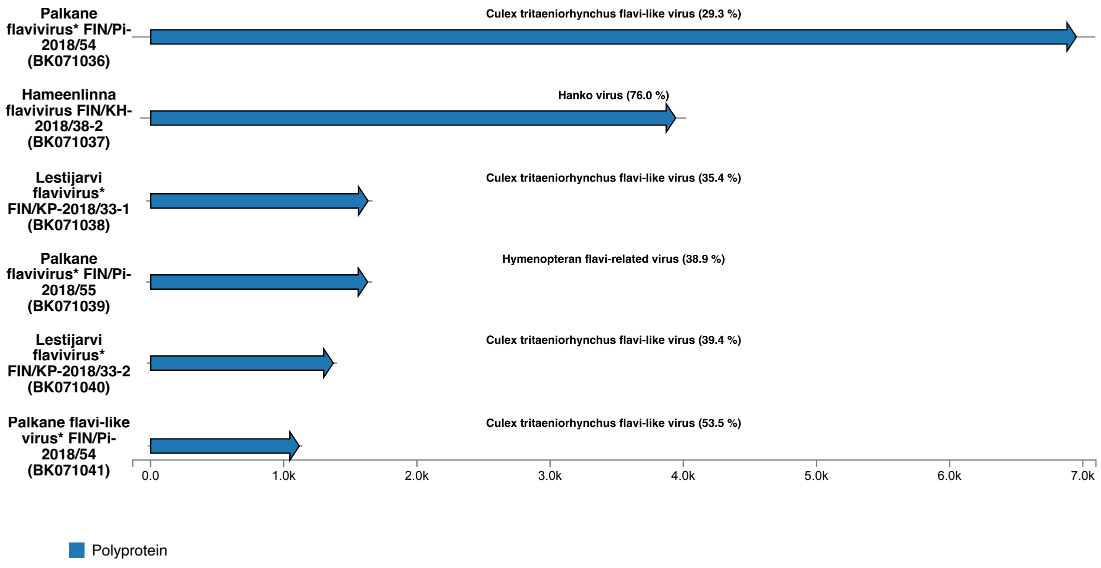


*Figure 5 ORF structure of novel Flavi-like sequences from mosquito samples. Labels at the center represent closest homologs in UniRef with identity values in parenthesis. *Provisional virus names, isolates and accession numbers.*

Bootstrapped ML tree was constructed for the six flavivirus-like sequences (≥ 1 kbp) as described in the Methods (Fig 6). *Hameenlinna flavivirus FIN/KH-2018/38-2* was identified as a new isolate of the previously published *Hameenlinna flavivirus FIN/KH-2018/38* (acc. ON955057). *Palkane flavi-like virus* (acc. BK071041) was placed as an outgroup to the *Placeda* (acc. MW434111) and *Culex tritaeniorhynchus flavi-like* (acc. LC514290) viruses.

Four novel sequences, including two isolates of *Lestijarvi flavivirus* (acc. BK071038 and BK071040) obtained from *Ochlerotatus diantaeus* and two isolates of *Palkane flavivirus* (acc. BK071036 and BK071039) from *O. communis*, formed a monophyletic group highly distant from previously published *Flaviviridae* and BLASTX homologs (Figure 6). Due to partial coverage of the polyprotein, the *Lestijarvi* and *Palkane* sequences were not characterised in details; however, this preliminary analysis indicates that these may represent isolates of novel flaviviruses infecting *O. diantaeus* and *O. communis* mosquitoes in Finland. Note that *Palkane flavi-like virus*, *Lestijarvi flavivirus*, and *Palkane flavivirus* are provisional names subject to revision.

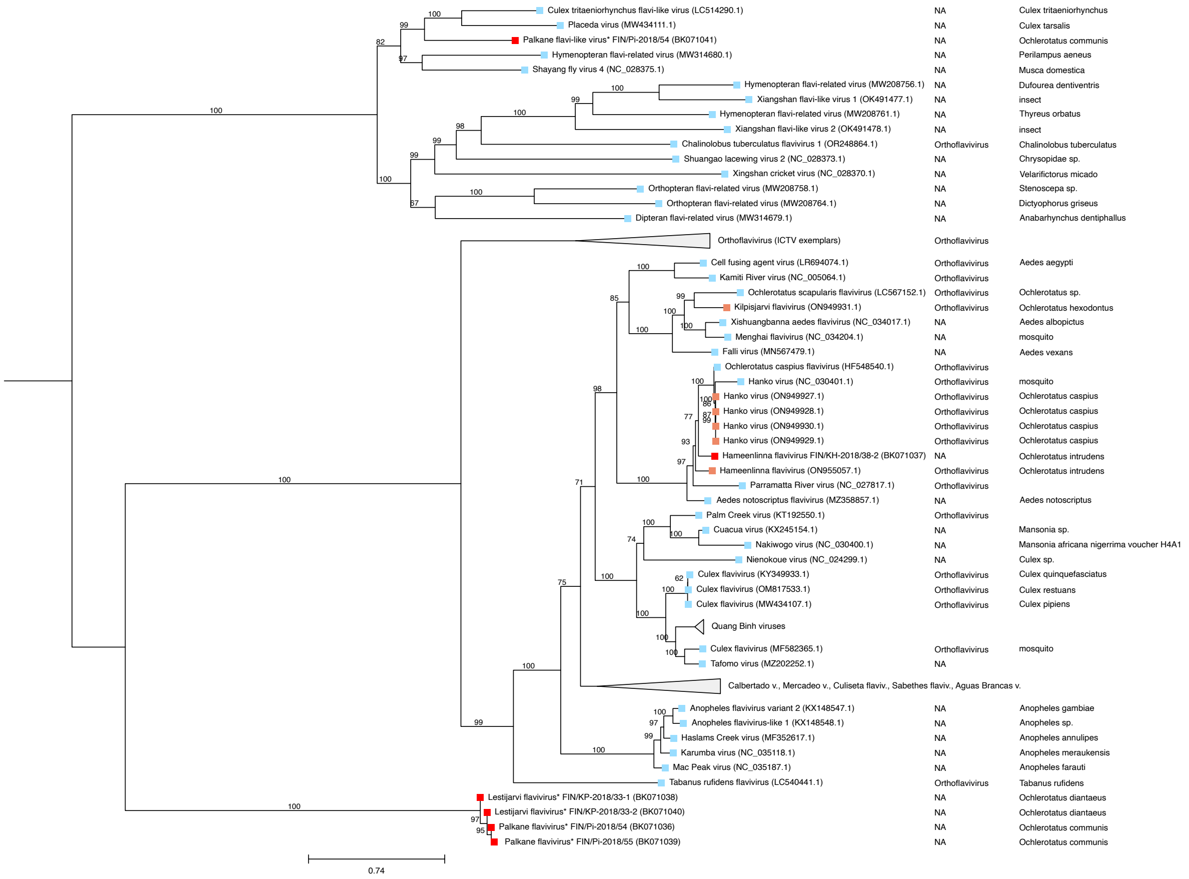


*Figure 6 Phylogeny of novel Flavi-like sequences from mosquito samples. Tip colours: novel sequences in red, homologs previously published for this data in light red, BLASTX homologs in blue, ICTV Orthoflavivirus exemplars in a collapsed clade in the upper part of the tree. Columns on the right denote genus and host species. Scale bar is in Mya. *Provisional virus names, isolates and accession numbers.*
